## Supplementary Information for "The hidden cost of receiving favors: A theory of indebtedness"

**This PDF file includes:**

#### SI Methods

##### Word classification task

An independent sample of participants was recruited (N = 80) to categorize the 100 words with the highest weight in the definitions of indebtedness collected in the large-scale online questionnaire in Study 1. They were instructed as following:

*In this experiment, you will see a list of words that people use in everyday helping and receiving helping situations. After seeing each word, please classify it into one of the following five categories according to your understanding:*

*1. Appraisal: if you think that the word is related to appraisals or evaluations on the benefits, costs, intentions or other factors in the situation, please classify the word into this category.*

*2. Emotion (Feeling): if you think that this word is related to or reflects certain emotions and feelings of an individual in the situation, please classify the word into this category.*

*3. Behavior: if you think the word is related to the behavior of an individual in the situation, please classify the word into this category.*

*4. Person: if you think this word is related to person, please classify the word into this category.*

*5. Other: if you think the word has nothing to do with any of the four categories above, please classify the word into this category.*

**Model comparisons**

***Model comparison for reciprocity decisions.*** First, we compared an alternative linear formulation of our model (Model 1.2) to the original non-linear version (Eq 1, Model 1.1).

***Model 1.2:***

$$U(D_B) = \theta_B * (\gamma_B - D_B) - (1 - \theta_B) * (\phi_B * \max(\omega_B * \gamma_B - D_B, 0) + (1 - \phi_B) * \max(E_B^{''} - D_B, 0))$$

To examine the necessities of both communal concern and obligation, we compared the full model (Model 1.1) to models that solely include term for communal concern (Model 1.3) or obligation (Model 1.4) besides the self-interest term.

***Model 1.3:***

$$U(D_B) = \theta_B * \frac{\gamma_B - D_B}{\gamma_B} - (1 - \theta_B) * (\frac{\omega_B * \gamma_B - D_B}{\gamma_B})^2$$

***Model 1.4:***

$$U(D_B) = \theta_B * \frac{\gamma_B - D_B}{\gamma_B} - (1 - \theta_B) * (\frac{E_B^{''} - D_B}{\gamma_B})^2$$

We further compared Model 1.1 to a model that independently parameterized the communal concern and obligation terms (Model 1.5), in which the ranges of parameters were defined as [0, 1].

***Model 1.5:***

$$U(D_B) = \theta_B * \frac{\gamma_B - D_B}{\gamma_B} - (1 - \theta_B) * (\phi_{B1} * (\frac{\omega_B * \gamma_B - D_B}{\gamma_B})^2 + \phi_{B2} * (\frac{E_B^{''} - D_B}{\gamma_B})^2)$$

We also compared Model 1.1 to a model that independently parameterized the self-interest, communal concern and obligation terms (Model 1.6), in which the ranges of parameters were defined as [0, 1].

***Model 1.6:***

$$U(D_B) = \theta_B * \frac{\gamma_B - D_B}{\gamma_B} - \phi_{B1} * (\frac{\omega_B * \gamma_B - D_B}{\gamma_B})^2 - \phi_{B2} * (\frac{E_B'' - D_B}{\gamma_B})^2$$

Another possibility was that participants did not make decisions based on feelings of communal concern and obligation, but instead decided how much to reciprocate simply according to the benefactor's cost. The closer the amount of reciprocity to the benefactor's cost, the larger the utility of this action. This is similar to standard models of reciprocity [1,2]. In the Repayment impossible and Repayment possible conditions, the weights on the benefactor's cost in the total utility were different, captured by parameters $\phi_{B1}$ and $\phi_{B2}$ respectively, resulting in the behavioral differences under different conditions (Model 1.7). In this model, $C$ is the condition-indicating coefficient: $C = 1$ represents Repayment impossible condition, and $C = 0$ represents the Repayment possible condition.

***Model 1.7****:*

$$U(D_B) = \theta_B * \pi_B + (1 - \theta_B) * (C * \phi_{B1} * U_{Cost} + (1 - C) * \phi_{B2} * U_{Cost})$$

$$U_{Cost} = -(\frac{D_A - D_B}{\gamma_B})^2$$

The third possibility is that participants made decisions according to inequity aversion [3,4], which assumes that participants care not only about self-interest, but also the payoff absolute difference between self and other (Model 1.8). The smaller the difference between self-payoff and the benefactor's payoff (or the closer the reciprocity is to 1/2 of their total payoff), the greater the utility of this action [4,5]. In the Repayment impossible and Repayment possible conditions, the weights on inequity were different, captured by parameters $\phi_1$ and $\phi_2$ respectively, resulting in the behavioral differences under different conditions.

***Model 1.8***:

$$U(D_B) = \theta_B * \pi_B + (1 - \theta_B) * (C * \phi_{B1} * U_{Inequity} + (1 - C) * \phi_{B2} * U_{Inequity})$$

$$U_{Inequity} = -(\frac{\gamma_B - D_B}{\gamma_A - D_A + D_B} - \frac{1}{2})^2$$

***Model comparison for help-acceptance decisions.*** Similar to reciprocity, to examine the necessities of both communal concern and obligation, we compared the full model (Model 2.1) to nested models that omitted the obligation (Model 2.2) and communal concern terms (Model 2.3). In all the models, *U(Reject)* was set to zero, because the participant's emotional responses would not change if the participant did not accept help.

***Model 2.2:***

$$U(Accept) = \theta_B * \frac{D_A * \mu}{\max(D_A * \mu)} + (1 - \theta_B) * \omega_B$$

***Model 2.3:***

$$U(Accept) = \theta_B * \frac{D_A * \mu}{\max(D_A * \mu)} - (1 - \theta_B) * \frac{E''_B}{\gamma_B}$$

We further compared the Model 2.1 to a model that had separate parameters for communal concern and obligation (Model 2.4), in which the ranges of $\theta$ and $\phi_2$ were defined as [0,1], and the ranges of $\phi_1$ was defined as [-1,1].

***Model 2.4***:

$$U(Accept) = \theta_B * \frac{D_A * \mu}{\max(D_A * \mu)} + (1 - \theta_B) * (\phi_{B1} * \omega_B - \phi_{B2} * \frac{E_B^{''}}{\gamma_B})$$

We also compared Model 2.1 to a model that independently parameterized the self-interest, communal concern and obligation terms (Model 2.5), in which the ranges of $\theta$ and $\phi_2$ were defined as [0,1], and the ranges of $\phi_1$ was defined as [-1,1].

***Model 2.5***:

$$U(Accept) = \theta_B * \frac{D_A * \mu}{\max(D_A * \mu)} + \phi_{B1} * \omega_B - \phi_{B2} * \frac{E_B^{''}}{\gamma_B}$$

### SI Results

#### Predicting behaviors using Communal and Obligation Factors

Because our computational model cannot differentiate between appraisals and feelings, we also report the results predicting behavior using the factors combining these ratings estimated from the exploratory factor analysis (EFA; Fig, 4D). We conducted linear mixed model for reciprocity by including the scores for Communal Factor and Obligation Factor extracted from EFA as fixed effects with by-participant random slopes for each fixed effect. Results demonstrated that both factors contributed significantly to reciprocity (Communal Factor: $\beta = 0.58 \pm 0.03$, $t = 17.20$, $p < 0.001$; Obligation Factor: $\beta = 0.20 \pm 0.02$, $t = 9.21$, $p < 0.001$). Similarly, both communal and obligation factors contributed significantly to the decisions of whether to accept help (Communal Factor: $\beta = -0.13 \pm 0.04$, $t = -17.55$, $p < 0.001$; Obligation Factor: $\beta = 0.06 \pm 0.03$, $t = 7.76$, $p < 0.001$). These results are consistent with separately predicting behavior from appraisals and emotions as reported in the main text.

#### Results of efficiency manipulation

In Study 2b, we further manipulated the participant's benefit from help by varying the exchange rate between the co-player's cost and participant's pain reduction (i.e., **Efficiency**, 0.5, 1, and 1.5) on the basis of Study 2a where Efficiency was 1. The higher the Efficiency, the more benefit the participant would obtain from each amount of the co-player's cost. Results are presented in Table S3-4. Specifically, first, in line with previous studies on gratitude showing that the beneficiary's benefit contributes positively to the feeling of gratitude [6-8], we found that the higher the Efficiency (i.e., the more benefit the participant obtained), the higher the participant's self-reported feeling of gratitude ($\beta = 0.05 \pm 0.02$, $t = 3.27$, $p < 0.001$). Similarly, participants' self-reported ratings of indebtedness were positively correlated with the size of Efficiency ($\beta = 0.04 \pm 0.01$, $t = 2.65$, $p = 0.008$). In contrary, the Efficiency did not contribute significantly to participants' rating of guilt and the sense of obligation (guilt: $\beta = 0.01 \pm 0.01$, $t = 0.74$, $p = 0.458$; obligation: $\beta = 0.00 \pm 0.02$, $t = -0.03$, $p = 0.975$) (Table S3-4).

To be noted, since the effects of Efficiency (or the benefit the beneficiary obtained from help) on beneficiary's emotions are not the focus of the current study, the range of Efficiency was relatively small in the current study (i.e., 0.5, 1, 1.5). Therefore, it is possible that the current manipulation was not efficient to capture some relatively extreme situations, which resulted in the current non-significant effects of Efficiency on guilt and the sense of obligation. For example, will a beneficiary feel less guilt when the benefactor's small cost has a larger effect, or feel more guilt when the benefactor's large cost has a small effect? Future specially designed studies are needed to explore how the efficiency of help influences the beneficiary's emotional responses.

#### Reference

1 Dufwenberg, M. & Kirchsteiger, G. A theory of sequential reciprocity. *Game. Econ. Behav.* **47**, 268-298, doi:10.1016/j.geb.2003.06.003 (2004).

2 Rabin, M. Incorporating fairness into game theory and economics. *Am. Econ. Rev.*, 1281-1302 (1993).

3 Bolton, G. E. & Ockenfels, A. ERC: A theory of equity, reciprocity, and competition. *Am. Econ. Rev.*, 166-193 (2000).

4 Fehr, E. & Schmidt, K. M. A theory of fairness, competition, and cooperation. *Q. J. Econ.* **114**, 817-868, doi:Doi 10.1162/003355399556151 (1999).

5 van Baar, J. M., Chang, L. J. & Sanfey, A. G. The computational and neural substrates of moral strategies in social decision-making. *Nat. Commun.* **10**, 1-14 (2019).

6 Tesser, A., Gatewood, R. & Driver, M. Some determinants of gratitude. *J. Pers. Soc. Psychol.* **9**, 233 (1968).

7 Yu, H., Gao, X., Zhou, Y. & Zhou, X. Decomposing gratitude: representation and integration of cognitive antecedents of gratitude in the brain. *J. Neurosci.*, 2944-2917 (2018).

8 Elfers, J. & Hlava, P. *The Spectrum of Gratitude Experience*. (Springer, 2016).

#### Supporting Tables

**Table S1. The contributions of guilt and obligation ratings to indebtedness rating**

| | **Model** | **Predictor** | **Df** | **AIC** | **Term** | **Beta** | **SE** | ***t*** | ***p*** |
|---|---|---|---|---|---|---|---|---|---|
| **Study 1** | Before controlling for benefactor's cost, participant's benefit and social distance | | | | | | | | |
| | ***Model 3 vs. Model 1**: $F$ = 1606.10, $p < 0.001$;* | | | | | | | | |
| | ***Model 3 vs. Model 2**: $F$ = 5.34, $p$ = 0.021; VIF = 1.22.* | | | | | | | | |
| | Model 1 | Obligation | 3 | 5412.8 | Obligation | 0.34 | 0.02 | 16.05 | < 0.001 |
| | Model 2 | Guilt | 3 | 4239.1 | Guilt | 0.71 | 0.02 | 45.41 | < 0.001 |
| | **Model 3** | Obligation | 4 | 4235.8 | Obligation | 0.40 | 0.02 | 2.31 | 0.021 |
| | | + Guilt | | | Guilt | 0.70 | 0.02 | 40.08 | < 0.001 |
| | After controlling for benefactor's cost, participant's benefit and social distance | | | | | | | | |
| | ***Model 3 vs. Model 1**: $F$ = 1005.70, $p < 0.001$;* | | | | | | | | |
| | ***Model 3 vs. Model 2**: $F$ = 3.53, $p$ = 0.060; VIF = 1.14.* | | | | | | | | |
| | Model 1 | Obligation | 3 | 19004.3 | Obligation | 0.42 | 0.03 | 15.87 | < 0.001 |
| | Model 2 | Guilt | 3 | 17913.3 | Guilt | 0.74 | 0.02 | 43.44 | < 0.001 |
| | **Model 3** | Obligation | 4 | 17907.6 | Obligation | 0.61 | 0.02 | 2.77 | 0.005 |
| | | + Guilt | | | Guilt | 0.71 | 0.02 | 38.26 | < 0.001 |
| **Study 2** | Before controlling for experimental variables | | | | | | | | |
| | ***Model 3 vs. Model 1**: $\chi^2$ = 1557.70, $df$ = 4, $p < 0.001$;* | | | | | | | | |
| | ***Model 3 vs. Model 2**: $\chi^2$ = 599.69, $df$ = 4, $p < 0.001$; VIF = 1.44.* | | | | | | | | |
| | Model 1 | Obligation | 6 | 6270.1 | Obligation | 0.35 | 0.05 | 6.51 | < 0.001 |
| | Model 2 | Guilt | 6 | 5312.0 | Guilt | 0.75 | 0.03 | 22.54 | < 0.001 |
| | **Model 3** | Obligation | 10 | 4720.3 | Obligation | 0.27 | 0.03 | 8.04 | < 0.001 |
| | | + Guilt | | | Guilt | 0.68 | 0.04 | 19.36 | < 0.001 |
| | After controlling for experimental variables | | | | | | | | |
| | ***Model 3 vs. Model 1**: $\chi^2$ = 245.44, $df$ = 4, $p < 0.001$;* | | | | | | | | |
| | ***Model 3 vs. Model 2**: $\chi^2$ = 97.871, $df$ = 4, $p < 0.001$; VIF = 1.03.* | | | | | | | | |
| | Model 1 | Obligation | 6 | 7744.6 | Obligation | 0.22 | 0.04 | 6.50 | < 0.001 |

| | Model | Predictor | Df | AIC | Term | Beta | SE | *t* | *p* |
|---|---|---|---|---|---|---|---|---|---|
| | Model 2 | Guilt | 6 | 7590.1 | Guilt | 0.34 | 0.03 | 11.30 | < 0.001 |
| | **Model 3** | Obligation | 10 | 7503.1 | Obligation | 0.17 | 0.03 | 10.23 | < 0.001 |
| | | + Guilt | | | Guilt | 0.30 | 0.03 | 6.24 | < 0.001 |
| **Study 3** | Before controlling for experimental variables | | | | | | | | |
| | ***Model 3 vs. Model 1**:* $\chi^2 = 462.43$, $df = 4$, $p < 0.001$; | | | | | | | | |
| | ***Model 3 vs. Model 2**:* $\chi^2 = 128.47$, $df = 4$, $p < 0.001$; VIF = 1.98. | | | | | | | | |
| | Model 1 | Obligation | 6 | 2319.2 | Obligation | 0.29 | 0.07 | 4.31 | < 0.001 |
| | Model 2 | Guilt | 6 | 1985.2 | Guilt | 0.63 | 0.04 | 15.83 | < 0.001 |
| | **Model 3** | Obligation | 10 | 1864.8 | Obligation | 0.20 | 0.05 | 3.82 | < 0.001 |
| | | + Guilt | | | Guilt | 0.62 | 0.06 | 13.64 | < 0.001 |
| | After controlling for experimental variables | | | | | | | | |
| | ***Model 3 vs. Model 1**:* $\chi^2 = 43.78$, $df = 4$, $p < 0.001$; | | | | | | | | |
| | ***Model 3 vs. Model 2**:* $\chi^2 = 17.25$, $df = 4$, $p = 0.002$; VIF = 1.00. | | | | | | | | |
| | Model 1 | Obligation | 6 | 2685.2 | Obligation | 0.17 | 0.04 | 4.36 | < 0.001 |
| | Model 2 | Guilt | 6 | 2658.8 | Guilt | 0.22 | 0.04 | 5.33 | < 0.001 |
| | **Model 3** | Obligation | 10 | 2649.6 | Obligation | 0.13 | 0.04 | 3.70 | < 0.001 |
| | | + Guilt | | | Guilt | 0.20 | 0.04 | 5.02 | < 0.001 |

**Table S2. Experimental designs for Study 2 and Study 3**

| Study | Study 2a | Study 2b | Study 3 |
|---|---|---|---|
| **Sample size** | 51 | 57 | 53 |
| **Conditions** | Benefactor knows: Repayment impossible vs. Repayment possible | Benefactor knows: Repayment impossible vs. Repayment possible | Benefactor knows: Repayment impossible vs. Repayment possible |
| **Cost** | 5, 7, 8, 9, 10, 11, 12,14, 15, 16, 18, 20 | 4, 8, 12, 16, 20 | 4, 6, 8, 10, 12, 14, 16, 18, 20 |
| **Efficiency manipulation** | 1 | 0.5, 1, 1.5 | 1 |
| **Dependent variables** | Reciprocity, Accept/Reject Help | Reciprocity, Accept/Reject Help | Reciprocity, |
| **Trial number of each cost-efficiency combination in each condition** | 1 | 1 | 3 |
| **Total trial number** | 48 | 56 | 54 |

**Table S3-1. The effects of the extra information about benefactor's intention, benefactor's cost, and efficiency on participants' emotional and behavioral responses (combining data of Studies 2a and 2b)**

| Dependent | Predictors | Beta | SE | *t* (*z*) | *p* |
|---|---|---|---|---|---|
| **Second-order belief** | Benefactor's cost | 0.42 | 0.02 | 20.84 | < 0.001 |
| | Extra information about benefactor's intention | 0.53 | 0.03 | 15.71 | < 0.001 |
| | Benefactor's cost×Extra information | 0.22 | 0.02 | 13.13 | < 0.001 |
| **Perceived care** | Benefactor's cost | 0.63 | 0.03 | 23.70 | < 0.001 |
| | Extra information about benefactor's intention | -0.31 | 0.02 | -13.89 | < 0.001 |
| | Benefactor's cost×Extra information | -0.08 | 0.01 | -6.64 | < 0.001 |
| **Gratitude** | Benefactor's cost | 0.55 | 0.03 | 19.36 | < 0.001 |
| | Extra information about benefactor's intention | -0.27 | 0.02 | -13.18 | < 0.001 |
| | Benefactor's cost×Extra information | -0.06 | 0.01 | -4.20 | < 0.001 |
| **Indebtedness** | Benefactor's cost | 0.52 | 0.03 | 20.24 | < 0.001 |
| | Extra information about benefactor's intention | -0.09 | 0.03 | -2.98 | 0.003 |
| | Benefactor's cost×Extra information | -0.01 | 0.01 | -0.72 | 0.474 |
| **Guilt** | Benefactor's cost | 0.41 | 0.02 | 17.04 | < 0.001 |
| | Extra information about benefactor's intention | -0.25 | 0.02 | -10.30 | < 0.001 |
| | Benefactor's cost×Extra information | -0.05 | 0.01 | -4.28 | < 0.001 |
| **Obligation** | Benefactor's cost | 0.22 | 0.03 | 7.71 | < 0.001 |
| | Extra information about benefactor's intention | 0.30 | 0.03 | 9.28 | < 0.001 |
| | Benefactor's cost×Extra information | 0.11 | 0.01 | 8.85 | < 0.001 |
| **Reciprocity** | Benefactor's cost | 0.63 | 0.02 | 25.60 | < 0.001 |
| | Extra information about benefactor's intention | -0.05 | 0.02 | -3.30 | 0.001 |
| | Benefactor's cost×Extra information | -0.03 | 0.01 | -2.99 | 0.003 |
| **Decision to reject help** | Benefactor's cost | -0.65 | 0.13 | -5.16 | < 0.001 |
| | Extra information about benefactor's intention | 0.27 | 0.08 | 3.64 | < 0.001 |
| | Benefactor's cost×Extra information | 0.07 | 0.07 | 1.08 | 0.279 |

**Table S3-2. The effects of the extra information about benefactor's intention and benefactor's cost on participants' emotional and behavioral responses (Study 2a)**

| Dependent variable | Predictors | Beta | SE | *t* (*z*) | *p* |
|---|---|---|---|---|---|
| **Second-order belief** | Benefactor's cost | 0.37 | 0.03 | 13.78 | < 0.001 |
| | Extra information about benefactor’s intention | 0.56 | 0.05 | 11.41 | < 0.001 |
| | Benefactor’s cost×Extra information | 0.18 | 0.02 | 7.38 | < 0.001 |
| **Perceived care** | Benefactor's cost | 0.51 | 0.04 | 14.57 | < 0.001 |
| | Extra information about benefactor’s intention | -0.38 | 0.04 | -9.80 | < 0.001 |
| | Benefactor’s cost×Extra information | -0.08 | 0.02 | -3.99 | < 0.001 |
| **Gratitude** | Benefactor's cost | 0.59 | 0.04 | 12.64 | < 0.001 |
| | Extra information about benefactor’s intention | -0.34 | 0.04 | -9.35 | < 0.001 |
| | Benefactor’s cost×Extra information | -0.05 | 0.02 | -2.54 | 0.014 |
| **Indebtedness** | Benefactor's cost | 0.33 | 0.03 | 11.26 | < 0.001 |
| | Extra information about benefactor’s intention | -0.27 | 0.04 | -6.46 | < 0.001 |
| | Benefactor’s cost×Extra information | -0.30 | 0.02 | -1.56 | 0.126 |
| **Guilt** | Benefactor’s cost | 0.43 | 0.03 | 12.31 | < 0.001 |
| | Extra information about benefactor’s intention | -0.07 | 0.06 | -1.10 | 0.277 |
| | Benefactor’s cost×Extra information | 0.00 | 0.02 | 0.08 | 0.935 |
| **Obligation** | Benefactor’s cost | 0.13 | 0.04 | 3.67 | < 0.001 |
| | Extra information about benefactor’s intention | 0.36 | 0.05 | 6.91 | < 0.001 |
| | Benefactor’s cost×Extra information | 0.17 | 0.02 | 6.22 | < 0.001 |
| **Reciprocity** | Benefactor’s cost | 0.64 | 0.03 | 22.92 | < 0.001 |
| | Extra information about benefactor’s intention | -0.05 | 0.03 | -1.66 | 0.103 |
| | Benefactor’s cost×Extra information | -0.03 | 0.02 | -1.47 | 0.147 |
| **Decision to reject help** | Benefactor’s cost | -0.37 | 0.21 | -1.70 | 0.088 |
| | Extra information about benefactor’s intention | 0.30 | 0.11 | 2.62 | 0.009 |
| | Benefactor’s cost×Extra information | 0.14 | 0.11 | 1.28 | 0.201 |

**Table S3-3. The effects of the extra information about benefactor's intention, and benefactor's cost on participants' emotional and behavioral responses (Study 2b)**

| Dependent variable | Predictors | Beta | SE | *t* (*z*) | *p* |
|---|---|---|---|---|---|
| **Second-order belief** | Benefactor's cost | 0.46 | 0.03 | 15.63 | < 0.001 |
| | Extra information about benefactor’s intention | 0.50 | 0.05 | 10.76 | < 0.001 |
| | Benefactor’s cost×Extra information | 0.25 | 0.02 | 10.91 | < 0.001 |
| **Perceived care** | Benefactor's cost | 0.72 | 0.04 | 19.49 | < 0.001 |
| | Extra information about benefactor’s intention | -0.25 | 0.02 | -10.55 | < 0.001 |
| | Benefactor’s cost×Extra information | -0.08 | 0.01 | -5.34 | < 0.001 |
| **Gratitude** | Benefactor's cost | 0.59 | 0.04 | 14.71 | < 0.001 |
| | Extra information about benefactor’s intention | -0.22 | 0.02 | -9.54 | < 0.001 |
| | Benefactor’s cost×Extra information | -0.05 | 0.02 | -3.27 | < 0.001 |
| **Indebtedness** | Benefactor's cost | 0.59 | 0.04 | 16.27 | < 0.001 |
| | Extra information about benefactor’s intention | -0.11 | 0.02 | -4.65 | < 0.001 |
| | Benefactor’s cost×Extra information | -0.02 | 0.01 | -1.40 | 0.164 |
| **Guilt** | Benefactor’s cost | 0.48 | 0.04 | 12.80 | < 0.001 |
| | Extra information about benefactor’s intention | -0.23 | 0.03 | -8.66 | < 0.001 |
| | Benefactor’s cost×Extra information | -0.07 | 0.01 | -4.74 | < 0.001 |
| **Obligation** | Benefactor’s cost | 0.30 | 0.04 | 7.27 | < 0.001 |
| | Extra information about benefactor’s intention | 0.25 | 0.04 | 6.25 | < 0.001 |
| | Benefactor’s cost×Extra information | 0.10 | 0.02 | 6.46 | < 0.001 |
| **Reciprocity** | Benefactor’s cost | 0.63 | 0.04 | 15.67 | < 0.001 |
| | Extra information about benefactor’s intention | -0.05 | 0.01 | -3.23 | < 0.001 |
| | Benefactor’s cost×Extra information | -0.04 | 0.01 | -2.92 | < 0.001 |
| **Decision to reject help** | Benefactor’s cost | -0.87 | 0.01 | -1262.41 | < 0.001 |
| | Extra information about benefactor’s intention | 0.24 | 0.01 | 351.12 | < 0.001 |
| | Benefactor’s cost×Extra information | 0.02 | 0.01 | 21.73 | < 0.001 |

**Table S3-4. The effects of the extra information about benefactor's intention, benefactor's cost, and efficiency on participants' emotional and behavioral responses (Study 2b)**

| Dependent variable | Predictors | Beta | SE | *t* (*z*) | *p* |
|---|---|---|---|---|---|
| **Second-order belief** | Benefactor's cost | 0.46 | 0.02 | 29.91 | <0.001 |
| | Extra information about benefactor’s intention | 0.51 | 0.02 | 33.12 | <0.001 |
| | Efficiency | 0.02 | 0.02 | 1.54 | 0.124 |
| | Benefactor’s cost×Extra information | 0.26 | 0.02 | 16.52 | <0.001 |
| | Efficiency×Benefactor’s cost | 0.01 | 0.02 | 0.42 | 0.678 |
| | Efficiency× Extra information | 0.01 | 0.02 | 0.89 | 0.376 |
| | Efficiency×Extra information ×Extra information | 0.02 | 0.02 | 1.36 | 0.174 |
| **Perceived care** | Benefactor's cost | 0.73 | 0.01 | 55.50 | <0.001 |
| | Extra information about benefactor’s intention | -0.25 | 0.01 | -19.37 | <0.001 |
| | Efficiency | 0.04 | 0.01 | 3.30 | 0.001 |
| | Benefactor’s cost×Extra information | -0.08 | 0.01 | -6.12 | <0.001 |
| | Efficiency×Benefactor’s cost | 0.02 | 0.01 | 1.71 | 0.087 |
| | Efficiency× Extra information | -0.01 | 0.01 | -0.72 | 0.469 |
| | Efficiency×Extra information ×Extra information | 0.00 | 0.01 | -0.25 | 0.803 |
| **Gratitude** | Benefactor's cost | 0.60 | 0.02 | 39.33 | <0.001 |
| | Extra information about benefactor’s intention | -0.22 | 0.02 | -14.96 | <0.001 |
| | Efficiency | 0.05 | 0.02 | 3.27 | 0.001 |
| | Benefactor’s cost×Extra information | -0.05 | 0.02 | -3.54 | <0.001 |
| | Efficiency×Benefactor’s cost | 0.00 | 0.02 | 0.01 | 0.989 |
| | Efficiency× Extra information | -0.01 | 0.02 | -0.86 | 0.388 |
| | Efficiency×Extra information ×Extra information | 0.01 | 0.02 | 0.40 | 0.688 |
| **Indebtedness** | Benefactor's cost | 0.60 | 0.01 | 40.90 | <0.001 |
| | Extra information about benefactor’s intention | -0.11 | 0.01 | -7.88 | <0.001 |
| | Efficiency | 0.04 | 0.01 | 2.65 | 0.008 |
| | Benefactor’s cost×Extra information | -0.03 | 0.01 | -1.79 | 0.074 |
| | Efficiency×Benefactor’s cost | 0.03 | 0.02 | 1.89 | 0.059 |
| | Efficiency× Extra information | -0.02 | 0.01 | -1.25 | 0.210 |
| | Efficiency×Extra information ×Extra information | -0.02 | 0.02 | -1.39 | 0.165 |
| **Guilt** | Benefactor's cost | 0.48 | 0.01 | 35.83 | <0.001 |

| | | | | | |
|---|---|---|---|---|---|
| | Extra information about benefactor’s intention | -0.23 | 0.01 | -17.01 | <0.001 |
| | Efficiency | 0.01 | 0.01 | 0.74 | 0.458 |
| | Benefactor’s cost×Extra information | -0.07 | 0.01 | -5.17 | <0.001 |
| | Efficiency×Benefactor’s cost | 0.01 | 0.01 | 0.57 | 0.567 |
| | Efficiency× Extra information | -0.02 | 0.01 | -1.32 | 0.188 |
| | Efficiency×Extra information ×Extra information | 0.00 | 0.01 | 0.22 | 0.828 |
| **Obligation** | Benefactor's cost | 0.30 | 0.02 | 19.60 | <0.001 |
| | Extra information about benefactor’s intention | 0.25 | 0.02 | 16.50 | <0.001 |
| | Efficiency | 0.00 | 0.02 | -0.03 | 0.975 |
| | Benefactor’s cost×Extra information | 0.10 | 0.02 | 6.43 | <0.001 |
| | Efficiency×Benefactor’s cost | 0.01 | 0.02 | 0.85 | 0.396 |
| | Efficiency× Extra information | -0.01 | 0.02 | -0.72 | 0.470 |
| | Efficiency×Extra information ×Extra information | -0.01 | 0.02 | -0.35 | 0.726 |
| **Reciprocity** | Benefactor's cost | 0.65 | 0.01 | 48.73 | <0.001 |
| | Extra information about benefactor’s intention | -0.05 | 0.01 | -4.00 | <0.001 |
| | Efficiency | 0.08 | 0.01 | 5.86 | <0.001 |
| | Benefactor’s cost×Extra information | -0.04 | 0.01 | -2.74 | 0.006 |
| | Efficiency×Benefactor’s cost | 0.02 | 0.01 | 1.69 | 0.090 |
| | Efficiency× Extra information | 0.00 | 0.01 | -0.08 | 0.940 |
| | Efficiency×Extra information ×Extra information | 0.01 | 0.01 | 0.63 | 0.528 |
| **Decision to reject help** | Benefactor's cost | -0.70 | 0.07 | -9.61 | <0.001 |
| | Extra information about benefactor’s intention | 0.27 | 0.07 | 3.92 | <0.001 |
| | Efficiency | -0.46 | 0.07 | -6.49 | <0.001 |
| | Benefactor’s cost×Extra information | 0.01 | 0.07 | 0.13 | 0.895 |
| | Efficiency×Benefactor’s cost | -0.11 | 0.07 | -1.51 | 0.131 |
| | Efficiency× Extra information | 0.05 | 0.07 | 0.67 | 0.500 |
| | Efficiency×Extra information ×Extra information | -0.04 | 0.07 | -0.53 | 0.594 |

Note: In the models of Table S3-4, we only included random intercepts for participants due to convergence issues with models that additionally included random slopes.

**Table S4. The effects of the extra information about benefactor's intention and benefactor's cost on participants' emotional and behavioral responses (fMRI study)**

| Dependent variable | Predictors | Beta | SE | *t* (*z*) | *p* |
|---|---|---|---|---|---|
| **Second-order belief** | Benefactor's cost | 0.44 | 0.02 | 22.20 | < 0.001 |
| | Extra information about benefactor's intention | 0.65 | 0.04 | 15.88 | < 0.001 |
| | Benefactor's cost×Extra information | 0.29 | 0.02 | 15.21 | < 0.001 |
| **Perceived care** | Benefactor's cost | 0.66 | 0.03 | 19.23 | < 0.001 |
| | Extra information about benefactor's intention | -0.29 | 0.03 | -10.00 | < 0.001 |
| | Benefactor's cost×Extra information | -0.09 | 0.02 | -4.16 | < 0.001 |
| **Gratitude** | Benefactor's cost | 0.66 | 0.03 | 20.45 | < 0.001 |
| | Extra information about benefactor's intention | -0.28 | 0.03 | -9.61 | < 0.001 |
| | Benefactor's cost×Extra information | -0.06 | 0.02 | -2.77 | 0.008 |
| **Indebtedness** | Benefactor's cost | 0.46 | 0.04 | 11.89 | < 0.001 |
| | Extra information about benefactor's intention | -0.26 | 0.04 | -6.64 | < 0.001 |
| | Benefactor's cost×Extra information | -0.09 | 0.02 | -4.05 | < 0.001 |
| **Guilt** | Benefactor's cost | 0.56 | 0.04 | 15.15 | < 0.001 |
| | Extra information about benefactor's intention | -0.11 | 0.03 | -3.45 | 0.001 |
| | Benefactor's cost×Extra information | 0.02 | 0.02 | 0.87 | 0.388 |
| **Obligation** | Benefactor's cost | 0.23 | 0.05 | 4.89 | < 0.001 |
| | Extra information about benefactor's intention | 0.29 | 0.04 | 6.67 | < 0.001 |
| | Benefactor's cost×Extra information | 0.17 | 0.02 | 7.53 | < 0.001 |
| **Reciprocity** | Benefactor's cost | 0.77 | 0.03 | 23.56 | < 0.001 |
| | Extra information about benefactor's intention | -0.07 | 0.02 | -4.21 | < 0.001 |
| | Benefactor's cost×Extra information | -0.02 | 0.01 | -2.65 | 0.009 |

**Table S5. Correlations between appraisals and emotions (within participant)**

| | | Second-order belief | Obligation | Indebtedness | Guilt | Gratitude | Perceived care |
|---|---|---|---|---|---|---|---|
| Second-order belief | average *r* | - | 0.47 | 0.28 | 0.09 | 0.13 | 0.13 |
| | *p* | - | < 0.001 | < 0.001 | 0.022 | < 0.001 | 0.001 |
| Obligation | average *r* | - | - | 0.27 | 0.10 | 0.04 | 0.05 |
| | *p* | - | - | < 0.001 | 0.032 | 0.275 | 0.175 |
| Indebtedness | average *r* | - | - | - | 0.66 | 0.55 | 0.67 |
| | *p* | - | - | - | < 0.001 | < 0.001 | < 0.001 |
| Guilt | average *r* | - | - | - | - | 0.64 | 0.75 |
| | *p* | - | - | - | - | < 0.001 | < 0.001 |
| Gratitude | average *r* | - | - | - | - | - | 0.80 |
| | *p* | - | - | - | - | - | < 0.001 |
| Perceived care | average *r* | - | - | - | - | - | - |
| | *p* | - | - | - | - | - | - |

**Table S6. Correlations between appraisals and emotions (between participant)**

| | | Second-order belief | Obligation | Indebtedness | Guilt | Gratitude | Perceived care |
|---|---|---|---|---|---|---|---|
| Second-order belief | average *r* | - | 0.36 | 0.25 | 0.27 | 0.00 | 0.17 |
| | *p* | - | < 0.001 | 0.009 | 0.005 | 0.996 | 0.086 |
| Obligation | average *r* | - | - | 0.44 | 0.42 | -0.06 | 0.03 |
| | *p* | - | - | < 0.001 | < 0.001 | 0.569 | 0.751 |
| Indebtedness | average *r* | - | - | - | 0.81 | 0.41 | 0.53 |
| | *p* | - | - | - | < 0.001 | < 0.001 | < 0.001 |
| Guilt | average *r* | - | - | - | - | 0.40 | 0.41 |
| | *p* | - | - | - | - | < 0.001 | < 0.001 |
| Gratitude | average *r* | - | - | - | - | - | 0.68 |
| | *p* | - | - | - | - | - | < 0.001 |
| Perceived care | average *r* | - | - | - | - | - | - |
| | *p* | - | - | - | - | - | - |

**Table S7. Model comparison for reciprocity decisions**

| | | Average Sum of Squared Error | | | Average AIC | | |
|---|---|---|---|---|---|---|---|
| Model | Model description | Study 2a | Study 2b | Combined | Study 2a | Study 2b | Combined |
| **Model 1.1** | Nonlinear version | 4331.00 | 3938.60 | 4123.90 | 117.51 | 130.49 | 124.36 |
| Model 1.2 | Linear version | 5550.93 | 7177.44 | 6409.36 | 125.22 | 148.06 | 137.27$^{***}$ |
| Model 1.3 | Only communal concern | 4773.74 | 5085.24 | 4938.14 | 117.16 | 136.07 | 127.14$^{*}$ |
| Model 1.4 | Only obligation | 40365.60 | 33791.45 | 36895.91 | 175.24 | 190.87 | 183.49$^{***}$ |
| Model 1.5 | Three separate parameters independently weighted communal concern and obligation | 4521.81 | 4134.60 | 4317.45 | 121.34 | 135.09 | 128.59$^{***}$ |
| Model 1.6 | Three separate parameters independently weighted greedy, communal concern and obligation | 4466.52 | 3840.01 | 4135.86 | 121.23 | 132.08 | 126.96* |
| Model 1.7 | Reciprocity according to benefactor's cost | 5475.37 | 5003.66 | 5226.41 | 124.01 | 133.75 | 129.15$^{***}$ |
| Model 1.8 | Inequity aversion model | 13726.42 | 10080.51 | 11802.19 | 150.92 | 160.58 | 156.02$^{***}$ |

Note: $^{*}$ Indicates AIC is significantly larger than that of Model 1.1 using Wilcoxon rank sum test. $^{*}p < 0.05$, $^{***}p < 0.001$.

**Table S8. Model comparison for help-acceptance decisions**

| | | Log Likelihood | | | Average AIC | | |
|---|---|---|---|---|---|---|---|
| Model | Model description | Study 2a | Study 2b | Combined | Study 2a | Study 2b | Combined |
| **Model 2.1** | Full model | -206.97 | -293.05 | -251.24 | 555.95 | 760.09 | 660.93 |
| Model 2.2 | Only communal concern | -202.92 | -303.73 | -254.77 | 549.85 | 775.47 | 665.88 |
| Model 2.3 | Only obligation | -237.20 | -339.40 | -289.76 | 570.40 | 790.80 | 683.75 |
| Model 2.4 | Three separate parameters independently weighted communal concern and obligation | -206.97 | -297.05 | -253.30 | 605.95 | 818.10 | 715.06$^{***}$ |
| Model 2.5 | Three separate parameters independently weighted greedy, communal concern and obligation | -208.69 | -293.17 | -253.14 | 607.56 | 821.33 | 717.50$^{***}$ |

Note: $^{*}$ Indicates AIC is significantly larger than that of Model 2.1 using Wilcoxon rank sum test. $^{***}p < 0.001$.

**Table S9. Parameter recovery for reciprocity decisions**

| Model | Model description | Study 2a | | Study 2b | | Total | |
|---|---|---|---|---|---|---|---|
| | | *r* ± *SE* | *p* | *r* ± *SE* | *p* | *r* ± *SE* | *p* |
| **Model 1.1** | Nonlinear version | .93±.07 | <0.001 | .94±.08 | <0.001 | .94±.07 | <0.001 |
| Model 1.2 | Linear version | .29±.20 | <0.001 | .36±.20 | <0.001 | .33±.20 | <0.001 |
| Model 1.3 | Only Communal Concern | .99±.02 | <0.001 | .90±.09 | <0.001 | .93±.06 | <0.001 |
| Model 1.4 | Only Obligation | .99±.02 | <0.001 | .98±.04 | <0.001 | .99±.03 | <0.001 |
| Model 1.5 | Three separate parameters independently weighted communal concern and obligation | .56±.28 | <0.001 | .65±.25 | <0.001 | .61±.27 | <0.001 |
| Model 1.6 | Three separate parameters independently weighted greedy, communal concern and obligation | .52±.28 | <0.001 | .70±.23 | <0.001 | .57±.27 | <0.001 |
| Model 1.7 | Reciprocity according to benefactor's cost | .80±.21 | <0.001 | .82±.19 | <0.001 | .81±.20 | <0.001 |
| Model 1.8 | Inequity aversion model | .76±.22 | <0.001 | .73±.24 | <0.001 | .74±.23 | <0.001 |

**Table S10. Parameter recovery for help-acceptance decisions**

| Model | Model description | Study 2a | | Study 2b | | Total | |
|---|---|---|---|---|---|---|---|
| | | *r* ± *SE* | *p* | *r* ± *SE* | *p* | *r* ± *SE* | *p* |
| **Model 2.1** | Full model | .45±.37 | <0.001 | .41±.38 | <0.001 | .43±.40 | <0.001 |
| Model 2.2 | Only Communal Concern | .73±.43 | <0.001 | .50±.48 | <0.001 | .62±.46 | <0.001 |
| Model 2.3 | Only Obligation | -.05±6.94 | 0.604 | .29±.67 | 0.002 | -.02±.4.80 | 0.738 |
| Model 2.4 | Three separate parameters independently weighted communal concern and obligation | .64±.32 | <0.001 | .62±.28 | <0.001 | .63±.30 | <0.001 |
| Model 2.5 | Three separate parameters independently weighted greedy, communal concern and obligation | .46±.32 | <0.001 | .43±.35 | <0.001 | .45±.34 | <0.001 |

**Table S11. Model estimated parameters for reciprocity decisions**

| | Study 2a | | Study 2b | | Combined | |
|---|---|---|---|---|---|---|
| Parameters | Mean | SE | Mean | SE | Mean | SE |
| $\theta$ | 0.06 | 0.01 | 0.10 | 0.02 | 0.08 | 0.01 |
| $W_{Communal}$ ($\phi$) | 0.83 | 0.03 | 0.75 | 0.02 | 0.79 | 0.02 |
| $W_{Obligation}$ ($1 - \phi$) | 0.17 | 0.03 | 0.25 | 0.02 | 0.21 | 0.02 |
| $\kappa$ | 0.21 | 0.03 | 0.41 | 0.04 | 0.32 | 0.01 |

**Table S12. Model estimated parameters for help-acceptance decisions**

| | Study 2a | | Study 2b | | Combined | |
|---|---|---|---|---|---|---|
| Parameters | Mean | SE | Mean | SE | Mean | SE |
| $\theta$ | 0.39 | 0.03 | 0.37 | 0.04 | 0.37 | 0.02 |
| $W_{Communal}$ ($\phi$) | -0.16 | 0.08 | 0.17 | 0.09 | 0.01 | 0.06 |
| $W_{Obligation}$ ($1 - \lvert\phi\rvert$) | 0.55 | 0.05 | 0.35 | 0.04 | 0.44 | 0.03 |
| $\kappa$ | 0.46 | 0.03 | 0.43 | 0.04 | 0.45 | 0.02 |

**Table S13. Results of whole-brain analysis of fMRI data**

| Regions | Hemisphere | *t* | Cluster size (voxels) | MNI coordinates | | |
|---|---|---|---|---|---|---|
| | | | | x | y | z |
| **Regions responded parametrically to the amount of reciprocity** | | | | | | |
| Left dlPFC | L | 5.93 | 209 | -45 | 5 | 29 |
| Right dlPFC | R | 4.91 | 138 | 45 | 11 | 35 |
| | | | | 39 | 8 | 38 |
| | R | 4.65 | 67 | 45 | 35 | 11 |
| Left IPL | L | 4.25 | 48 | -54 | -40 | 53 |
| Right IPL | R | 4.80 | 130 | 51 | -28 | 47 |
| Precuneus-MOG-ITG | R | 6.59 | 958 | 51 | -52 | -13 |
| MOG | L | 5.34 | 637 | -30 | -67 | 29 |
| ITG | L | 5.30 | 479 | -45 | -61 | -13 |
| Cerebellum | L | 38 | 3.99 | -27 | -61 | -34 |
| Cerebellum | R | 45 | 4.55 | 6 | -31 | -22 |
| **Regions responded parametrically to communal concern ($\omega_B$)** | | | | | | |
| vmPFC | - | 4.19 | 41 | 0 | 35 | -22 |
| aINS | L | 4.19 | 23 | -24 | 11 | -19 |
| Left dlPFC | L | 4.35 | 79 | -48 | 20 | 26 |
| | L | 4.60 | 75 | -24 | 29 | 56 |
| Right dlPFC | R | 5.24 | 251 | 45 | 11 | 38 |
| Precuneus | R | 4.46 | 683 | 3 | -46 | 38 |
| ITG | R | 4.85 | 128 | 48 | -46 | -16 |
| ITG | L | 4.64 | 198 | -54 | -76 | -7 |
| MOG | L | 4.89 | 490 | -30 | -76 | 32 |
| Calcarine | R | 4.29 | 61 | 9 | -91 | 5 |
| **Regions identified in parametric contrast for obligation ($E_B''$)** | | | | | | |
| dmPFC | L | 4.39 | 31 | -9 | 44 | 41 |
| Left TPJ | L | 3.86 | 42 | -57 | -61 | 26 |

Note: vmPFC = ventromedial prefrontal cortex; aINS = anterior insula; IPL = inferior parietal lobule; dlPFC = dorsolateral prefrontal cortex; ITG = Inferior temporal gyrus; dmPFC = dorsomedial prefrontal lobe; dACC = dorsal anterior cingulate gyrus; MOG = middle occipital gyrus. All brain maps were thresholded using cluster correction FWE $p < 0.05$ with a cluster-forming threshold of $p < 0.001$.

**Table S14. The meaning of symbols for variables in the computational model**

| Variable | Meaning |
|---|---|
| $D$ | Decision over choice space, e.g., the benefactor's cost, the beneficiary's amount of reciprocity and the beneficiary's decision of accepting help |
| $\theta$ | Greed sensitivity |
| $\phi$ | Mixture weight of $U_{Communal}$ and $U_{Obligation}$ |
| $\gamma$ | Endowment size |
| $\pi$ | Self interest |
| $\mu$ | The efficiency of help |
| $E''$ | Second-order belief of how much the benefactor expects |
| $\omega$ | Perceived care |
| $\kappa$ | The influence of second-order belief ($E''$) on perceived care ($\omega$) (higher indicates lower perceived care) |
| $\lambda$ | Inverse temperature parameter |
| $n$ | Total number of trials |
| $t$ | Trial number |

#### Supporting Figures

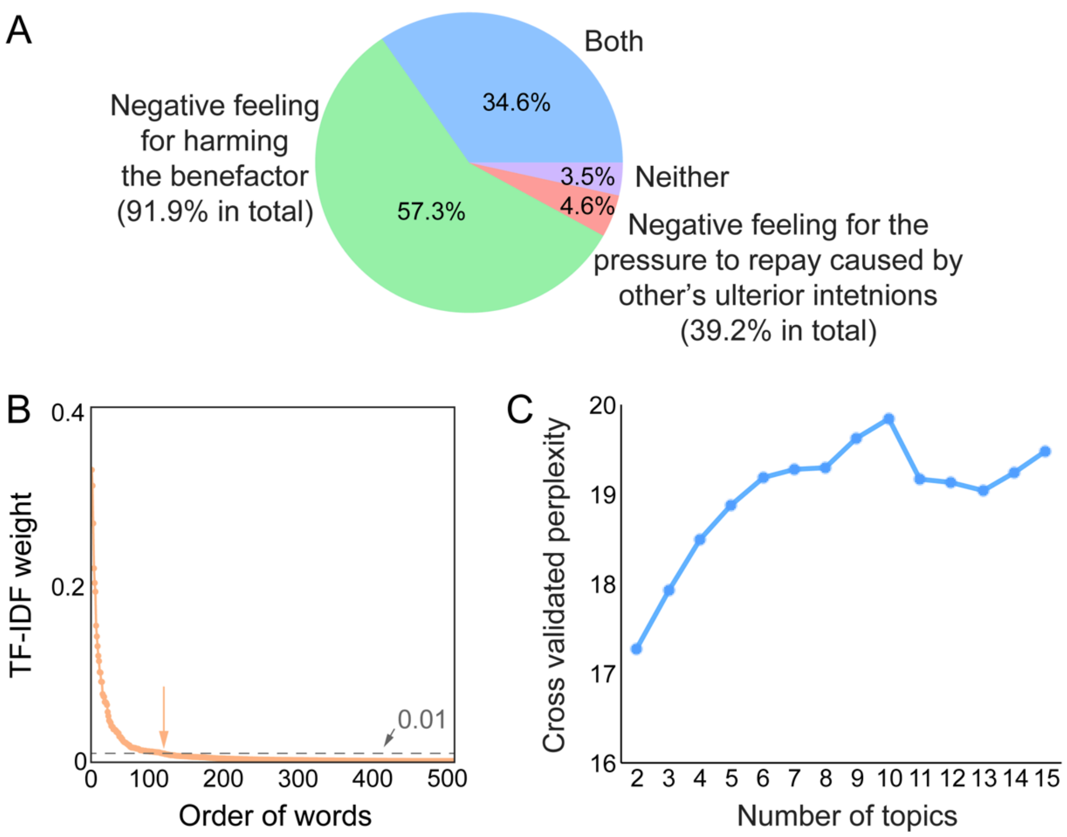

**Fig. S1 Definition of indebtedness. (A)** The frequency of choosing each option in the question "In daily life, what do you think is/ are the source(s) of indebtedness?" While 57.3% of participants indicated the negative feeling for harming the benefactor as the single source of indebtedness, 4.6% of participants indicated the negative feeling for the pressure of repayment caused by other's ulterior intentions as the single source of indebtedness. 34.6% of participants indicated both types of negative emotions contributing to indebtedness, and 3.5% of participants indicated neither of them as the source of indebtedness. **(B)** Emotional words were extracted from the 100 words with the highest weight/frequency in the definitions of indebtedness based on the annotation by an independent sample of participants (N = 80). Note, words beyond these 100 had TF-IDF weights < 0.01, indicating that the words included in the current analysis explained vast majority of variance in the definition of indebtedness. **(C)** We selected the best number of topics by comparing the models with topic numbers ranging from 2 to 15 using 5-folds cross validation. Model goodness of fit was assessed using perplexity, with lower perplexity denoting a better probabilistic model. We found that the two-topic solution performed the best.

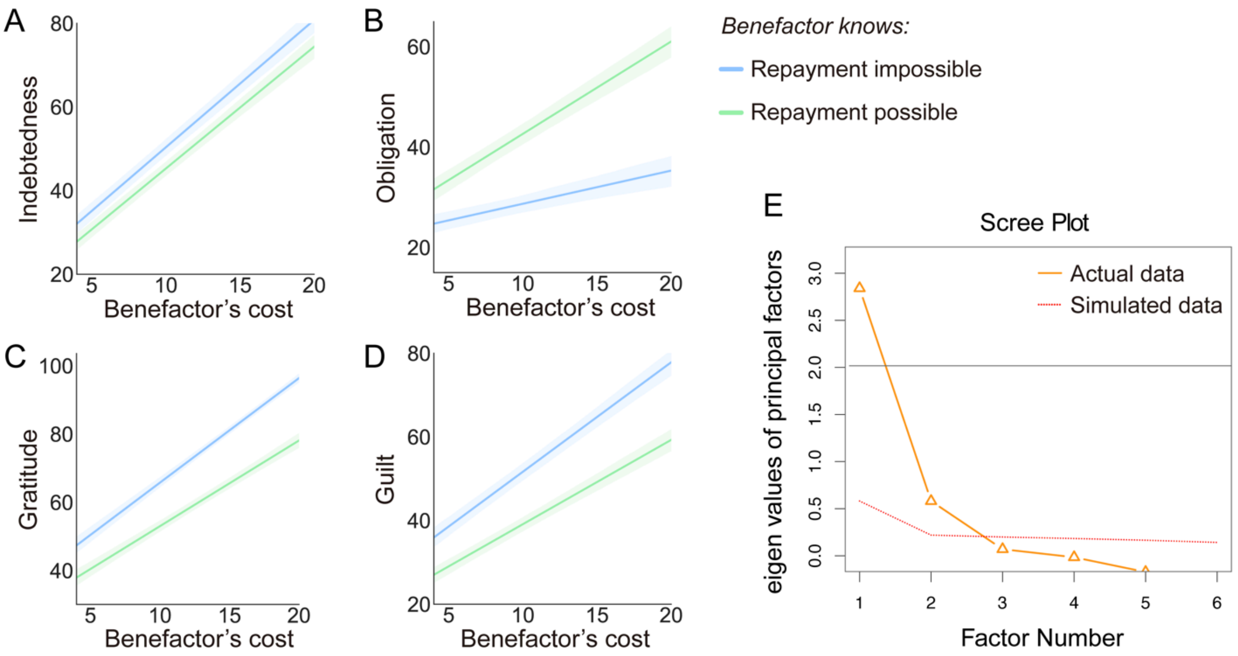

**Fig. S2 (A-D)** Participant's ratings on indebtedness, obligation, gratitude and guilt plotted as functions of the extra information about benefactor's intention and benefactor's cost in Study 2. **(E)** To determine the number of factors to retain in the exploratory factor analysis (EFA) in Study 2, the correlation matrix between appraisals and emotions was submitted to a parallel analysis [14]. Parallel analysis performed a principal factor decomposition of the data matrix and compared it to a principal factor decomposition of a randomized data matrix. This analysis yielded factors whose eigenvalues (magnitudes) were greater in the observed data relative to the randomized data. The nScree function was used to return an analysis of the number of factors to retain. The result pointed to a two-factor solution except for the acceleration factor.

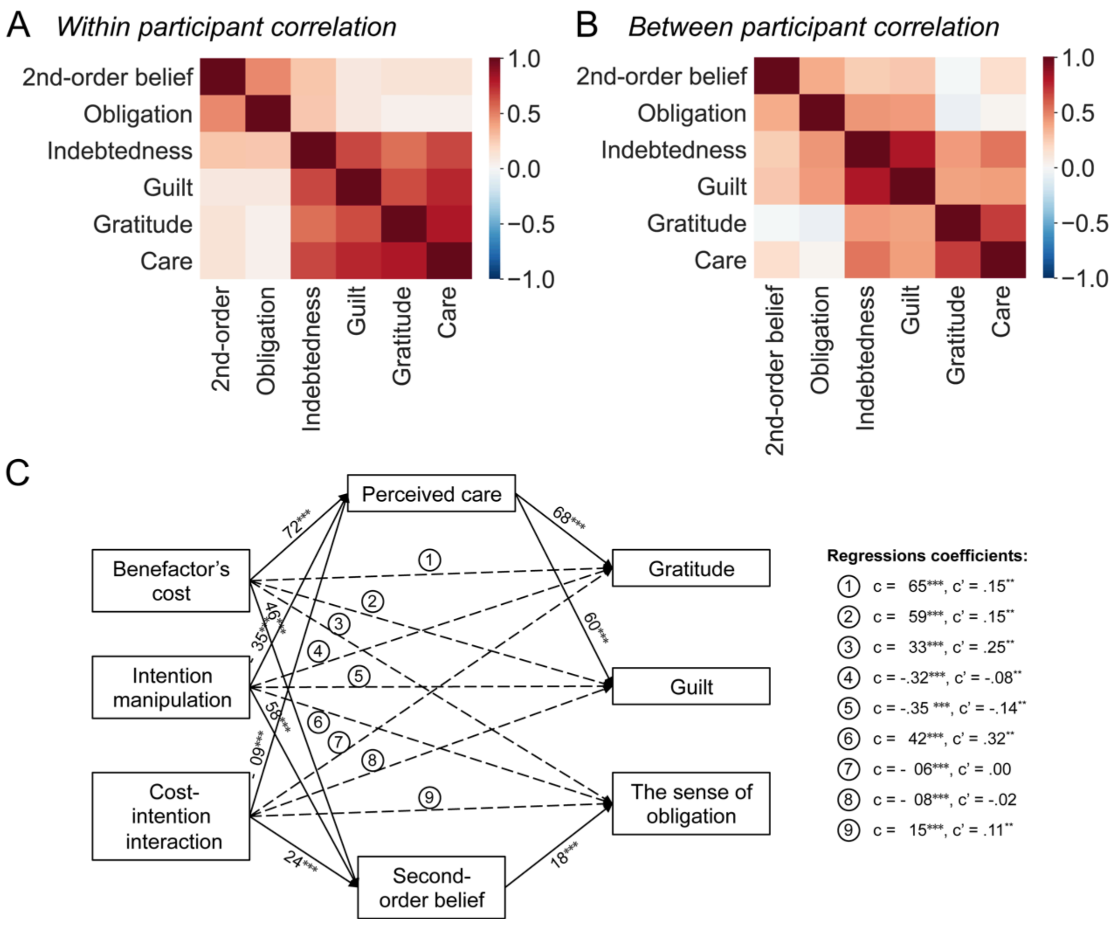

**Fig. S3 Relationships between appraisals and emotions. (A-B)** Correlation matrix between participant's appraisal and emotion ratings built based on within participant (A) and between participant (B) variances, respectively. **(C)** Mediation analysis showed that second-order beliefs and perceived care appraisals differentially mediated the effects of the experimental manipulations on emotional responses (total indirect effect = 0.59±0.04, $Z$ = 14.49, $p$ < 0.001; Fig. 4E and Fig. S3C). Second-order beliefs mediated the effects of the experimental manipulations on the sense of obligation (Indirect effect = 0.22±0.03, $Z$ = 7.18, $p$ < 0.001), while perceived care mediated the effects of the experimental manipulations on guilt (Indirect effect = 0.17±0.01, $Z$ = 13.23, $p$ < 0.001) and gratitude (Indirect effect = 0.19±0.01, $Z$ = 13.72, $p$ < 0.001).

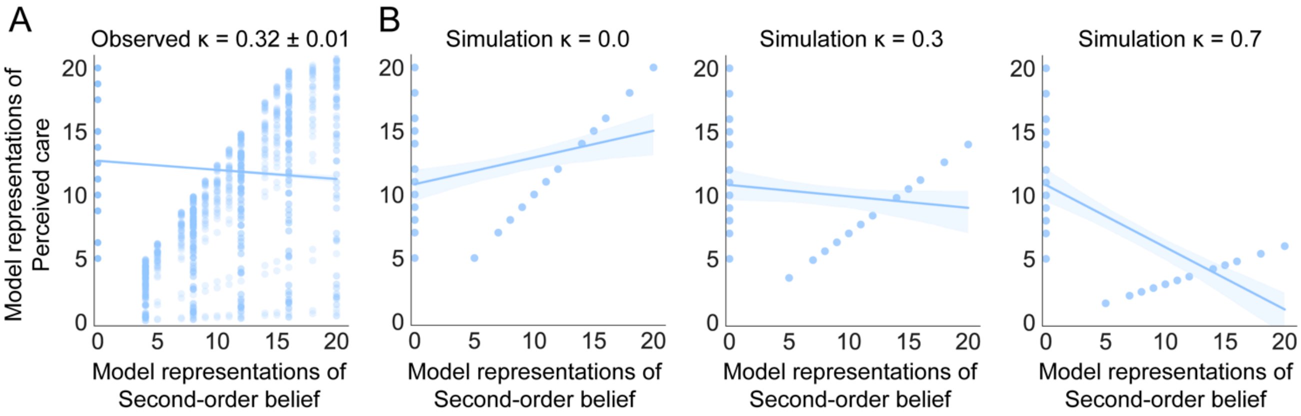

**Fig. S4** Relationships between model representations of second-order belief $E_B''$ and perceived care $\omega_B$ in the current study (A) and the simulated data with different levels of $\kappa$ (B).

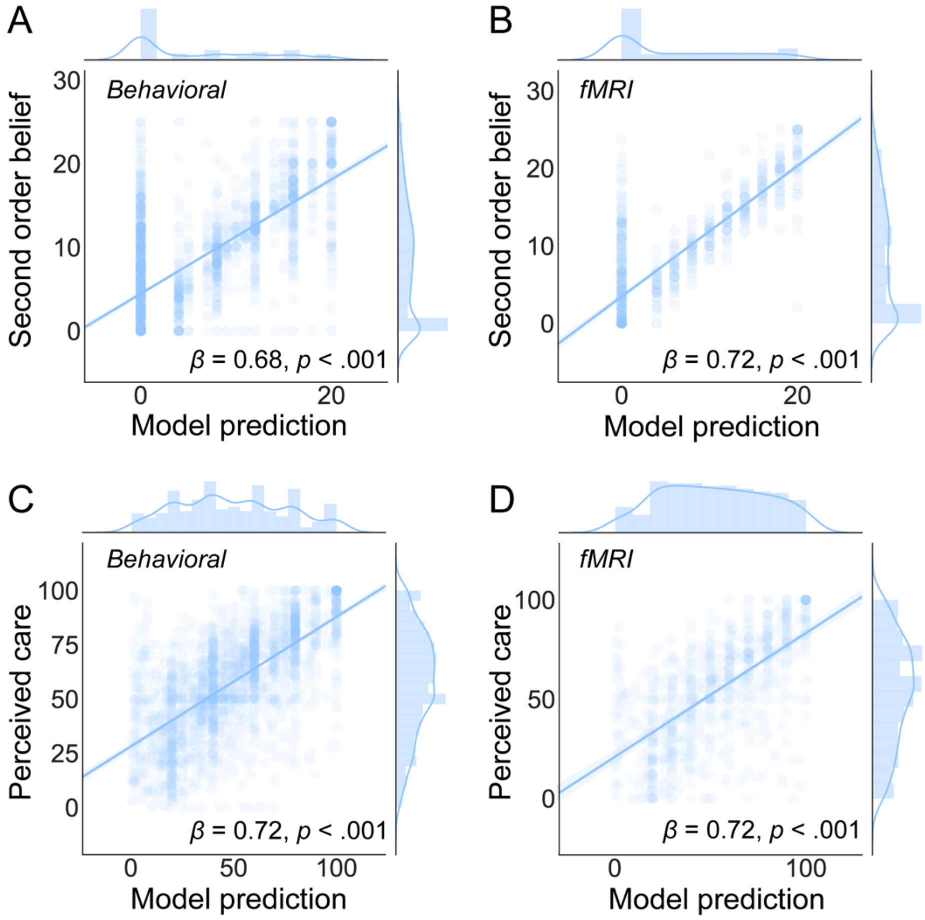

**Fig. S5 Model predictions for appraisals on benefactor's intentions.** A regression analysis showed that the reciprocity model's representations of $E_B''$ and $\omega_B$ were associated with trial-to-trial variations in self-reported values of second-order belief of the benefactor's expectation for repayment ($\beta_{behavioral}$ = 0.68 ± 0.03 (mean ± SE), $t$ = 21.48, $p < 0.001$; $\beta_{fMRI}$ = 0.84 ± 0.04, $t$ = 21.89, $p < 0.001$, A and B) and perceived care ($\beta_{behavioral}$ = 0.64 ± 0.02, $t$ = 26.76, $p < 0.001$; $\beta_{fMRI}$ = 0.70 ± 0.04, $t$ = 17.78, $p <$ 0.001, C and D), which provided further validation that the model representations were reflecting the intended psychological processes.

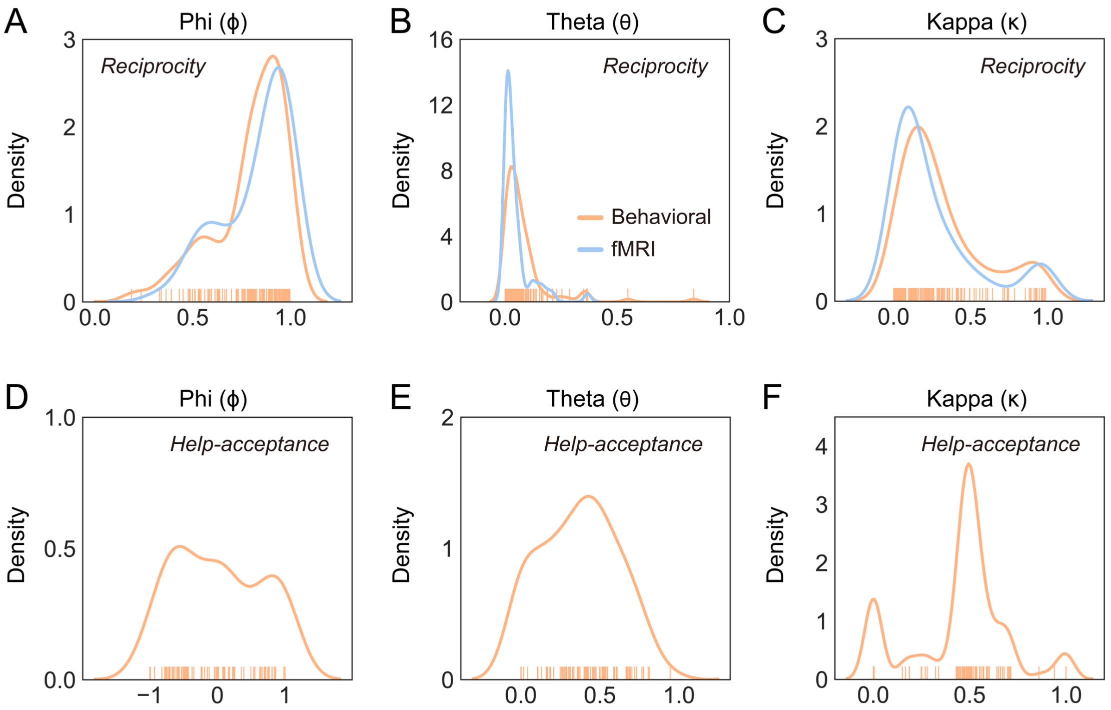

**Fig. S6 Distributions of model parameters. (A - C)** Distributions of parameters for reciprocity decisions in behavioral and fMRI studies. **(D - F)** Distributions of parameters for help-acceptance decisions in the behavioral study.

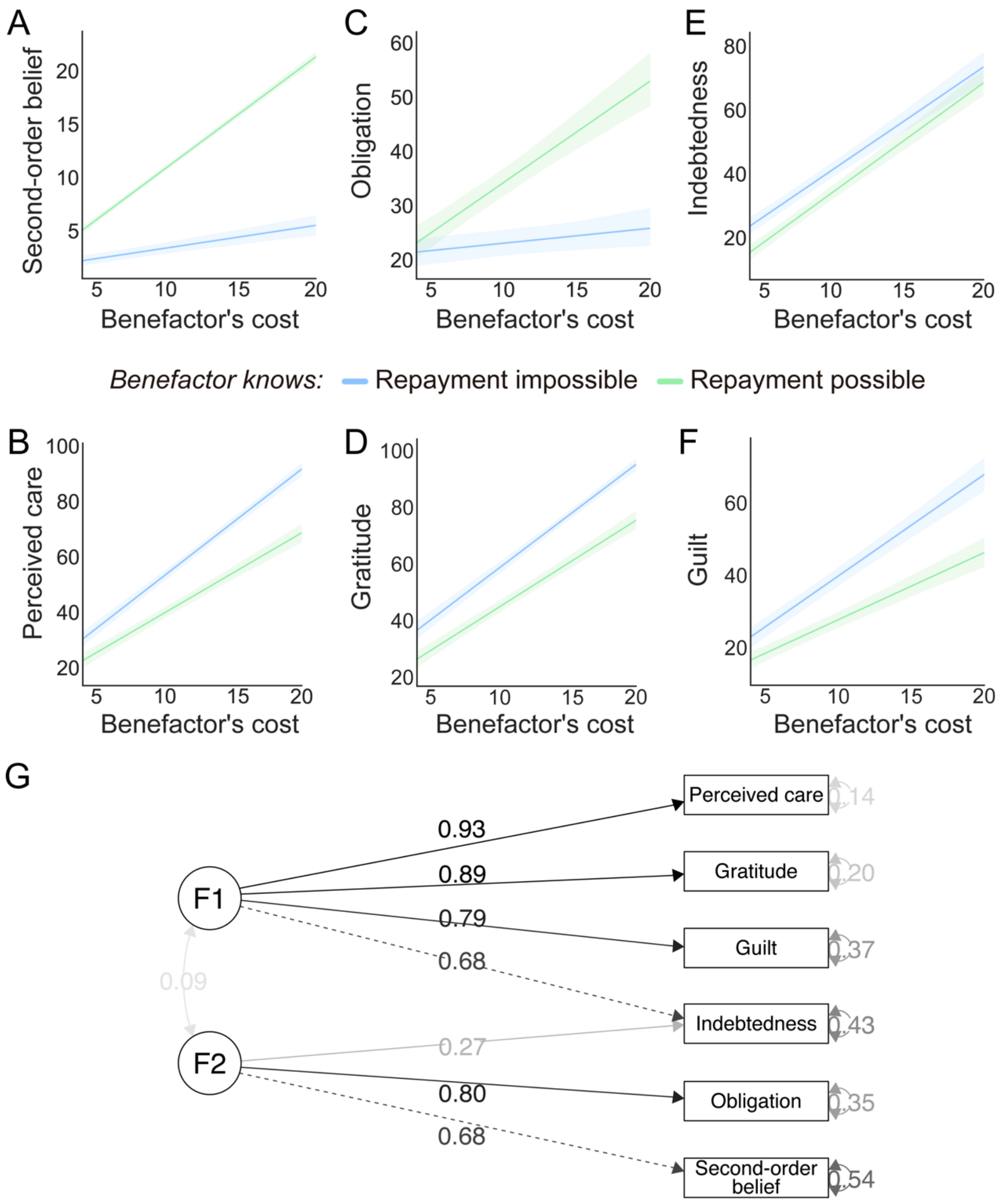

**Fig. S7 Appraisal and emotional responses in the fMRI study (Study 3) replicated that in the behavioral studies.** Compared with the Repayment impossible condition, the participant's second-order belief of the benefactor's expectation for repay (A), the sense of obligation (C) increased, while the participant's perceived care (B), gratitude (D) and guilt (F) decreased in Repayment possible condition. As the increase of benefactor's cost, the sizes of these effects increased (i.e., significant interaction effects between the extra information about benefactor's intention and benefactor's cost). Participants reported feeling of indebtedness in both conditions, but slightly more in the Repayment impossible compared to the Repayment possible condition (E). (G) We conducted confirmatory factor analysis (CFA) in Study 3 to test the two-factor model (Fig. 4E) built by Study 2 in an independent sample. Results showed that the fitness of this two-factor model is appropriate (RSMEA = 0.079, SRMR = 0.019, CFI = 0.986, TLI = 0.970).

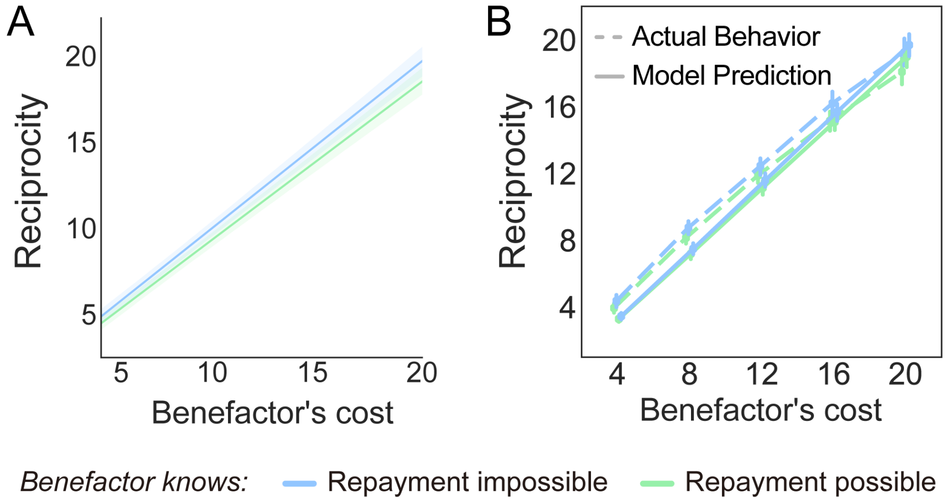

**Fig. S8 The results of reciprocity in the fMRI study (Study 3) replicates that in the behavioral studies (Study 2). (A)** Participants reciprocated more as the benefactor's increased. This effect was slightly enhanced in the Repayment impossible relative to the Repayment possible condition. **(B)** Our computational model accurately captured the patterns of participants' reciprocity after receiving help, $r^2 = 0.95$, $p <$ 0.001.

**Appendices S1. Online questionnaire for Study 1**

(Note: The questions included in final analysis are labeled in bold.)

*Welcome to participate in this questionnaire survey! In the questionnaire, please recall your real life events, and answer the corresponding questions. After answering all the questions, please write a short story about each event. Your story may be used as material for future study. If your story is selected, we will contact you through your contact information and pay you 25 yuan for story authorization. When using the story, we will keep your personal information strictly confidential. Please fill in the answer sheet sincerely and carefully.*

*To ensure the quality of the data, please check the following box, and promise that you will have at least 15 minutes to conduct the survey and answer each question sincerely. Thank you for your cooperation!*

○ I guarantee that I have at least 15 minutes to fill out the questionnaire and answer each question sincerely.

**Part 1**

Have you received any help in the past one year? [single choice]

○ Yes (Continue)  ○ No (skip Part 1)

Please think carefully about an event in which you received help from others that impressed you the most in the past one year and happened recently.

1. What is the time of this event? (Please select the option that matches the occurrence of the event and is closest to today)

○Within one week ○Within one month,

○Within three months ○Within half a year ○Within one year

2. In this event, did you actively seek help or passively accept help from others? [single choice]

○Actively seek  ○Passive acceptance

**3. How was your relationship with this benefactor before receiving help? (0 is very unfamiliar, 100 is very familiar)**

4. To what extent were you willing to accept this help? (0 is not willing to accept, 100

very willing to accept)

5. Who was the person that helped you? [single choice]

○Parent ○Sibling ○Spouse/boyfriend/girlfriend ○Other relative ○Friend

○Classmate/Colleague ○Teacher ○Neighbor ○Stranger

○Other (such as service personnel, public servants, etc.. Please fill in the benefactor's specific occupation) ________________

**6. How helpful was the help? (0 is useless, 100 is very helpful)**

**7. How much was the benefactor's cost in this help? (0 is not at all, 100 is very big)**

8. Before receiving the help, how likely did you think the benefactor would help? (0 is completely impossible, 100 is pretty sure)

**9. How grateful did you feel about the benefactor's help? (0 is not at all, 100 is very strong)**

**10. How indebted did you feel about the benefactor's help? (0 is not at all, 100 is very strong)**

**11. How guilty did you feel about the benefactor's help? (0 is not at all, 100 is very strong)**

**12. How much were you afraid of the benefactor's expectation for repay? (0 is not at all, 100 is very strong)**

13. How much pressure did you feel to reciprocate in the future? (0 is not at all, 100 is very strong)

**14. To what extent did you think you needed to reciprocate? (0 is not at all, 100 is very strong)**

**15. To what extent do you think the benefactor expected you to repay? (0 is not at all, 100 is very strong)**

16. Compared with the benefit you obtained from the help, how much did you think you needed to reciprocate to the benefactor? (0-50 means less than your benefit, 50 means equal to your benefit, 50-100 means more than your benefit)

17. Compared with the benefactor's cost, how much did you think you needed to reciprocate to the benefactor? (0-50 means less than the benefactor's cost, 50 means equal to the benefactor's cost, 50-100 means more than the benefactor's cost)

18. Compared with the benefit you obtained from the help, how much did you think the benefactor needed you to reciprocate? (0-50 means less than your benefit, 50 means

equal to your benefit, 50-100 means more than your benefit)

19. Compared with the benefactor's cost, how much did you think the benefactor needed you to reciprocate? (0-50 means less than the benefactor's cost, 50 means equal to the benefactor's cost, 50-100 means more than the benefactor's cost)

20. Did the benefactor propose a clear request for reciprocity? [single choice]

○ Yes, please briefly explain the details _________________ ○ No

**21. To what extent did you think the benefactor cared about your welfare when helping you? (0 is not at all, 100 is very strong)**

22. To what extent did you think the benefactor cared about his/her own interests when helping you? (0 is not at all, 100 is very strong)

23. To what extent did you want to reciprocate? (0 is not at all, 100 is very strong)

24. To what extent did you want to repay the favor immediately? (0 is not at all, 100 is very strong)

25. In what way did you want to reciprocate? [multiple choice]

□ Monetary reciprocity □ Help each other

□ Oral thanks □ Establish cooperative relationship

□ Make friends with him/her □ Gifts □ Other _________________

26. Have you reciprocated in some way? [multiple choice]

□ Monetary reciprocity □ Help each other

□ Oral thanks □ Establish cooperative relationship

□ Make friends with him/her □ Gifts □ Other _________________

27. To what extent were you willing to interact or get to know each other further? (0 is not at all, 100 is very strong)

28. Please describe the event in detail in the form of a short story.

(If your story is selected as the material for our experiment, you will receive a story authorization fee of 25 yuan)

**Part 2**

Have you rejected any help in the past one-year? [single choice]

○ Yes (Continue) ○ No (skip Part 2)

Please think carefully about an event when you received help from others that impressed you most in the past one year and happened recently.

1. What is the time of this event? (Please select the option that matches the occurrence of the event and is closest to today)

○Within one week ○Within one month,

○Within three months ○Within half a year ○Within one year

2. In this event, did the benefactor actively provide the offer of help or did someone suggest the benefactor to give you help? [single choice]

○Actively provided the offer of help ○Someone suggested the benefactor to help

**3. How was your relationship with this benefactor before this event? (0 is very unfamiliar, 100 is very familiar)**

4. To what extent did you want to reject the offer of help? (0 is not at all, 100 is very strong)

5. Who was the person that offered to help you? [single choice]

○Parent ○Sibling ○Spouse/boyfriend/girlfriend ○Other relative ○Friend

○Classmate/Colleague ○Teacher ○Neighbor ○Stranger

○Other (such as service personnel, public servants, etc. Please fill in the benefactor's specific occupation) _________________

**6. Imagine if you have accepted the help, how helpful would the help be? (0 is useless, 100 is very helpful)**

**7. Imagine if you have accepted the help, how much would the benefactor's cost be in this help? (0 is not at all, 100 is very big)**

8. Before receiving the help, how likely did you think the benefactor would help? (0 is completely impossible, 100 is pretty sure)

**9. Imagine if you have accepted the help, how grateful would you feel about the benefactor's help? (0 is not at all, 100 is very strong)**

**10. Imagine if you have accepted the help, how indebted would you feel about the benefactor's help? (0 is not at all, 100 is very strong)**

**11. Imagine if you have accepted the help, how guilty would you feel about the benefactor's help? (0 is not at all, 100 is very strong)**

**12. Imagine if you have accepted the help, how much you were afraid of the**

**benefactor's expectation for repay? (0 is not at all, 100 is very strong)**

13. Imagine if you have accepted the help, how much pressure would you feel to reciprocate in the future? (0 is not at all, 100 is very strong)

**14. Imagine if you have accepted the help, to what extent did you think you needed to reciprocate? (0 is not at all, 100 is very strong)**

**15. Imagine if you have accepted the help, to what extent would you think the benefactor expected you to reciprocate? (0 is not at all, 100 is very strong)**

16. Imagine if you have accepted the help, compared with the benefit you obtained from the help, how much would you think you needed to reciprocate to the benefactor? (0-50 means less than your benefit, 50 means equal to your benefit, 50-100 means more than your benefit)

17. Imagine if you have accepted the help, compared with the benefactor's cost, how much would you think you needed to reciprocate to the benefactor? (0-50 means less than the benefactor's cost, 50 means equal to the benefactor's cost, 50-100 means more than the benefactor's cost)

18. Imagine if you have accepted the help, compared with the benefit you obtained from the help, how much would you think the benefactor needed you to reciprocate? (0-50 means less than your benefit, 50 means equal to your benefit, 50-100 means more than your benefit)

19. Imagine if you have accepted the help, compared with the benefactor's cost, how much would you think the benefactor needed you to reciprocate? (0-50 means less than the benefactor's cost, 50 means equal to the benefactor's cost, 50-100 means more than the benefactor's cost)

20. Did the benefactor ask for repayment before helping you? [single choice]

○Yes, please briefly explain the details _________________ ○No

**21. To what extent did you think the benefactor cared about your welfare when he/she offered to help you? (0 is not at all, 100 is very strong)**

22. To what extent did you think the benefactor cared about his/her own interests when he/she offered to help you? (0 is not at all, 100 is very strong)

23. Imagine if you have accepted the help, to what extent would you want to reciprocate? (0 is not at all, 100 is very strong)

24. Imagine if you have accepted the help, to what extent did you want to repay the favor immediately? (0 is not at all, 100 is very strong)

25. Imagine if you have accepted the help, in what way did you want to reciprocate? [multiple choice]

□ Monetary reciprocity □ Help each other

□ Oral thanks □ Establish cooperative relationship

□ Make friends with him/her □ Gifts □ Other

27. What was/were your reason(s) for refusing the offer? [multiple choice]

□ Thought the benefactor's purpose not pure

□ The anticipatory repayment was too much

□ Limit your freedom

□ Feeling your self-esteem was hurt

□ The benefit from the help was little

□ The benefactor's cost was too much

28. Please describe the event in detail in the form of a short story.

(If your story is selected as the material for our experiment, you will receive a story authorization fee of 25 yuan)

**• In the context of helping and receiving help, what is your definition of gratitude?**

**• In the context of helping and receiving help, what is your definition of indebtedness?**

**• In daily life, what do you think is/are the source(s) of indebtedness? (Single choice, the order of the first two options was counterbalanced among participants)**

**□ Negative feeling for harming the benefactor/for cost that the benefactor has paid for helping you**

**☐ Negative feeling for the pressure to repay caused by other's ulterior intentions (e.g., Expectation for repay)**

**☐ Both of the above**

**☐ Neither of the above**

**Appendices S2. Classification for words in the definition of indebtedness**

| Classification | Word | English word | Weight | Frequency to be Classified in each level (%) | | | | |
|---|---|---|---|---|---|---|---|---|
| | | | | Appraisal | Emotion | Behavior | Person | Other |
| Appraisal | 损失 | Loss | 0.075 | 45.0 | 7.5 | 36.3 | 0.0 | 11.3 |
| | 代价 | Cost | 0.033 | 41.3 | 5.0 | 17.5 | 2.5 | 33.8 |
| | 不好 | Bad | 0.014 | 55.0 | 30.0 | 0.0 | 1.3 | 13.8 |
| | 受损 | Harm | 0.012 | 45.0 | 6.3 | 45.0 | 0.0 | 3.8 |
| | 很大 | Great | 0.012 | 51.3 | 5.0 | 2.5 | 2.5 | 38.8 |
| | 不必要 | Unnecessary | 0.010 | 46.3 | 12.5 | 5.0 | 1.3 | 35.0 |
| Emotion | 愧疚 | Guilt | 0.269 | 1.3 | 97.5 | 0.0 | 0.0 | 1.3 |
| | 内疚 | Guilt | 0.192 | 0.0 | 98.8 | 0.0 | 1.3 | 0.0 |
| | 亏欠 | Feel indebted | 0.154 | 25.0 | 46.3 | 26.3 | 0.0 | 2.5 |
| | 感觉 | Feel | 0.120 | 10.0 | 66.3 | 15.0 | 0.0 | 8.8 |
| | 感到 | Feel | 0.102 | 5.0 | 66.3 | 20.0 | 0.0 | 8.8 |
| | 觉得 | Feel | 0.074 | 15.0 | 53.8 | 17.5 | 0.0 | 13.8 |
| | 对不起 | Feel sorry | 0.068 | 8.8 | 62.5 | 15.0 | 0.0 | 13.8 |
| | 想要 | Want to | 0.052 | 6.3 | 56.3 | 32.5 | 1.3 | 3.8 |
| | 不安 | Uneasy | 0.047 | 0.0 | 97.5 | 1.3 | 0.0 | 1.3 |
| | 麻烦 | Trouble | 0.041 | 26.3 | 36.3 | 21.3 | 2.5 | 13.8 |
| | 难受 | Uncomfortable | 0.040 | 0.0 | 98.8 | 0.0 | 1.3 | 0.0 |
| | 负罪感 | Guilt | 0.034 | 3.8 | 93.8 | 0.0 | 1.3 | 1.3 |
| | 自责 | Guilt | 0.033 | 5.0 | 85.0 | 8.8 | 1.3 | 0.0 |
| | 过意不去 | Feel sorry | 0.026 | 2.5 | 95.0 | 0.0 | 1.3 | 1.3 |
| | 有愧 | Guilt | 0.023 | 1.3 | 95.0 | 2.5 | 0.0 | 1.3 |
| | 感激 | Gratitude | 0.022 | 1.3 | 86.3 | 12.5 | 0.0 | 0.0 |
| | 不好意思 | Feel sorry | 0.020 | 1.3 | 91.3 | 2.5 | 3.8 | 1.3 |
| | 抱歉 | Feel sorry | 0.017 | 2.5 | 87.5 | 8.8 | 1.3 | 0.0 |
| | 不舒服 | Uncomfortable | 0.016 | 5.0 | 92.5 | 0.0 | 1.3 | 1.3 |
| | 心里 | In the heart | 0.013 | 3.8 | 41.3 | 25.0 | 3.8 | 26.3 |
| | 压力 | Pressure | 0.013 | 10.0 | 72.5 | 3.8 | 1.3 | 12.5 |
| | 情感 | Emotion | 0.013 | 7.5 | 70.0 | 0.0 | 0.0 | 22.5 |
| | 内疚感 | Guilt | 0.013 | 1.3 | 96.3 | 0.0 | 2.5 | 0.0 |
| | 负担 | Burden | 0.012 | 16.3 | 37.5 | 33.8 | 0.0 | 12.5 |
| | 痛苦 | Painful | 0.011 | 1.3 | 95.0 | 3.8 | 0.0 | 0.0 |
| | 强烈 | Strong | 0.010 | 20.0 | 57.5 | 2.5 | 1.3 | 18.8 |
| | 希望 | Want to | 0.010 | 16.3 | 56.3 | 17.5 | 0.0 | 10.0 |
| | 歉疚 | Guilt | 0.010 | 1.3 | 96.3 | 1.3 | 1.3 | 0.0 |
| Behavior | 帮助 | Help | 0.312 | 3.8 | 0.0 | 93.8 | 2.5 | 0.0 |
| | 伤害 | Harm | 0.101 | 15.0 | 11.3 | 73.8 | 0.0 | 0.0 |
| | 付出 | Cost | 0.091 | 10.0 | 3.8 | 82.5 | 1.3 | 2.5 |
| | 负债 | Be in debt | 0.090 | 22.5 | 13.8 | 46.3 | 2.5 | 15.0 |
| | 回报 | Repay | 0.068 | 20.0 | 3.8 | 66.3 | 1.3 | 8.8 |

| Classification | Word | English word | Weight | Frequency to be Classified in each level (%) | | | | |
|---|---|---|---|---|---|---|---|---|
| | | | | Appraisal | Emotion | Behavior | Person | Other |
| | 造成 | Cause | 0.067 | 15.0 | 0.0 | 78.8 | 0.0 | 6.3 |
| | 损害 | Harm | 0.065 | 22.5 | 5.0 | 68.8 | 0.0 | 3.8 |
| | 受到 | Receive | 0.046 | 7.5 | 17.5 | 45.0 | 0.0 | 30.0 |
| | 接受 | Receive | 0.044 | 3.8 | 10.0 | 82.5 | 1.3 | 2.5 |
| | 产生 | Generate | 0.037 | 7.5 | 3.8 | 57.5 | 1.3 | 30.0 |
| | 补偿 | Compensate | 0.037 | 12.5 | 3.8 | 83.8 | 0.0 | 0.0 |
| | 牺牲 | Sacrifice | 0.037 | 15.0 | 2.5 | 75.0 | 1.3 | 6.3 |
| | 偿还 | Repay | 0.034 | 16.3 | 2.5 | 77.5 | 1.3 | 2.5 |
| | 回馈 | Repay | 0.022 | 13.8 | 1.3 | 78.8 | 2.5 | 3.8 |
| | 带来 | Bring | 0.019 | 6.3 | 3.8 | 77.5 | 1.3 | 11.3 |
| | 收到 | Receive | 0.017 | 2.5 | 2.5 | 88.8 | 1.3 | 5.0 |
| | 需要 | Need | 0.017 | 21.3 | 17.5 | 32.5 | 1.3 | 27.5 |
| | 影响 | Influence | 0.016 | 26.3 | 11.3 | 51.3 | 0.0 | 11.3 |
| | 弥补 | Compensate | 0.016 | 5.0 | 10.0 | 83.8 | 1.3 | 0.0 |
| | 行为 | Behavior | 0.015 | 7.5 | 1.3 | 68.8 | 1.3 | 21.3 |
| | 给予 | Give | 0.013 | 3.8 | 0.0 | 93.8 | 0.0 | 2.5 |
| | 报答 | Repay | 0.012 | 8.8 | 7.5 | 81.3 | 1.3 | 1.3 |
| | 得到 | Receive | 0.012 | 6.3 | 2.5 | 82.5 | 1.3 | 7.5 |
| | 付出代价 | Pay the price | 0.012 | 17.5 | 2.5 | 70.0 | 1.3 | 8.8 |
| | 我会 | I will | 0.012 | 16.3 | 7.5 | 47.5 | 5.0 | 23.8 |
| | 做错 | Wrongdoings | 0.012 | 28.8 | 6.3 | 57.5 | 2.5 | 5.0 |
| | 做错事 | Wrongdoings | 0.011 | 17.5 | 5.0 | 72.5 | 0.0 | 5.0 |
| | 失去 | Loss | 0.011 | 15.0 | 10.0 | 67.5 | 0.0 | 7.5 |
| | 导致 | Lead to | 0.011 | 18.8 | 2.5 | 57.5 | 1.3 | 20.0 |
| Person | 别人 | Other | 0.329 | 1.3 | 0.0 | 1.3 | 96.3 | 1.3 |
| | 他人 | Other | 0.218 | 0.0 | 0.0 | 2.5 | 97.5 | 0.0 |
| | 自己 | Self | 0.202 | 0.0 | 0.0 | 0.0 | 98.8 | 1.3 |
| | 对方 | Other | 0.142 | 2.5 | 0.0 | 0.0 | 96.3 | 1.3 |
| | 帮助者 | Benefactor | 0.036 | 2.5 | 1.3 | 2.5 | 88.8 | 5.0 |
| | 其他人 | Other | 0.026 | 1.3 | 0.0 | 2.5 | 95.0 | 1.3 |
| | 自身 | Self | 0.016 | 2.5 | 3.8 | 0.0 | 83.8 | 10.0 |
| Other | 利益 | Benefit | 0.131 | 36.3 | 7.5 | 6.3 | 2.5 | 47.5 |
| | 内心 | In the heart | 0.114 | 3.8 | 43.8 | 0.0 | 2.5 | 50.0 |
| | 因为 | Because | 0.077 | 18.8 | 3.8 | 5.0 | 1.3 | 71.3 |
| | 心存 | In the heart | 0.057 | 6.3 | 36.3 | 2.5 | 1.3 | 53.8 |
| | 人情 | Favor | 0.040 | 17.5 | 20.0 | 7.5 | 2.5 | 52.5 |
| | 东西 | Things | 0.029 | 7.5 | 1.3 | 2.5 | 8.8 | 80.0 |
| | 心理 | Psychological | 0.029 | 7.5 | 38.8 | 1.3 | 2.5 | 50.0 |
| | 事情 | Things | 0.028 | 5.0 | 0.0 | 8.8 | 5.0 | 81.3 |
| | 或者 | Or | 0.024 | 7.5 | 0.0 | 1.3 | 1.3 | 90.0 |

| Classification | Word | English word | Weight | Frequency to be Classified in each level (%) | | | | |
|---|---|---|---|---|---|---|---|---|
| | | | | Appraisal | Emotion | Behavior | Person | Other |
| | 为了 | In order to | 0.023 | 18.8 | 2.5 | 21.3 | 1.3 | 56.3 |
| | 一种 | A kind of | 0.021 | 15.0 | 0.0 | 2.5 | 2.5 | 80.0 |
| | 没有 | No | 0.020 | 25.0 | 0.0 | 2.5 | 1.3 | 71.3 |
| | 某件事 | Something | 0.019 | 3.8 | 0.0 | 6.3 | 15.0 | 75.0 |
| | 有所 | Somewhat | 0.017 | 16.3 | 8.8 | 7.5 | 0.0 | 67.5 |
| | 对于 | For | 0.016 | 21.3 | 2.5 | 10.0 | 2.5 | 63.8 |
| | 一些 | Some | 0.015 | 11.3 | 0.0 | 3.8 | 6.3 | 78.8 |
| | 什么 | What | 0.015 | 7.5 | 2.5 | 1.3 | 2.5 | 86.3 |
| | 应该 | Should | 0.015 | 30.0 | 16.3 | 10.0 | 1.3 | 42.5 |
| | 程度 | Extent | 0.014 | 33.8 | 2.5 | 6.3 | 5.0 | 52.5 |
| | 由于 | Because | 0.013 | 23.8 | 1.3 | 5.0 | 1.3 | 68.8 |
| | 原因 | Reason | 0.013 | 25.0 | 3.8 | 3.8 | 0.0 | 67.5 |
| | 某事 | Something | 0.012 | 5.0 | 0.0 | 5.0 | 11.3 | 78.8 |
| | 某些 | Some | 0.012 | 7.5 | 0.0 | 5.0 | 7.5 | 80.0 |
| | 一定 | Certainly | 0.012 | 27.5 | 7.5 | 7.5 | 2.5 | 55.0 |
| | 是否 | Whether | 0.012 | 37.5 | 1.3 | 2.5 | 1.3 | 57.5 |
| | 感是 | Feel | 0.012 | 10.0 | 28.8 | 3.8 | 0.0 | 57.5 |
| | 心中 | In the heart | 0.011 | 3.8 | 40.0 | 2.5 | 3.8 | 50.0 |
| | 无法 | Cannot | 0.011 | 25.0 | 11.3 | 6.3 | 0.0 | 57.5 |
| | 道德 | Moral | 0.011 | 25.0 | 16.3 | 11.3 | 2.5 | 45.0 |
| | 责任 | Responsibility | 0.010 | 25.0 | 23.8 | 15.0 | 3.8 | 32.5 |
